# Cohesin drives chromatin scanning during the RAD51-mediated homology search

**DOI:** 10.1101/2025.02.10.637451

**Authors:** Alberto Marin-Gonzalez, Adam T. Rybczynski, Namrata M. Nilavar, Daniel Nguyen, Violetta Karwacki-Neisius, Andrew G. Li, Roger S. Zou, Franklin J. Avilés-Vázquez, Masato T. Kanemaki, Ralph Scully, Taekjip Ha

**Affiliations:** Howard Hughes Medical Institute and Program in Cellular and Molecular Medicine, Boston Children’s Hospital, Boston, MA, USA; Department of Pediatrics, Harvard Medical School, Boston, MA, USA; Department of Biology, Johns Hopkins University, Baltimore, MD, USA; Department of Medicine, Beth Israel Deaconess Medical Center and Harvard Medical School, Boston, MA, USA; Department of Medicine, Massachusetts General Hospital, Boston, MA, USA; Department of Biophysics and Biophysical Chemistry, Johns Hopkins University, Baltimore, MD, USA; Department of Chromosome Science, National Institute of Genetics, Mishima, Japan; Graduate Institute for Advanced Studies, SOKENDAI, Mishima, Japan; Department of Biological Science, Graduate School of Science, The University of Tokyo, Tokyo, Japan

## Abstract

Cohesin folds genomes into chromatin loops, whose roles are under debate. We report that double strand breaks (DSB) induce *de novo* formation of chromatin loops, with the break positioned at the loop base. These loops form only in S/G2 phases and occur during repair via homologous recombination (HR), concomitant with DNA end resection and RAD51 assembly. RAD51 showed two-tiered accumulation around DSBs, with a broad (~Mb) domain arising from the homology search. This domain is regulated by cohesin unloader, is constrained by TAD boundaries, and it overlaps with chromatin regions reeled through the break-anchored loop, suggesting that loop extrusion regulates the homology search. Indeed, depletion of NIPBL results in reduced HR, and this effect is more pronounced when the HR donor is far (~100 kb) from the break. Our data indicates that loop-extruding cohesin promotes the mammalian homology search by facilitating break-chromatin interactions within the damaged TAD.

**One-Sentence Summary:** High spatiotemporal resolution analysis of double strand beak repair in 3D genome revealed the role of cohesin-driven loop extrusion in the homology search.

## INTRODUCTION

At the scale of hundreds of kilobases to a few megabases, mammalian genomes are organized into topologically associating domains (TADs) and chromatin loops (*1, 2*). These structures are dependent on the activity of the cohesin complex and are likely formed via a loop extrusion mechanism, whereby cohesin binds to chromatin and extrudes a loop until a convergent CTCF is found (*3–7*). Genomic loci that fall within the same TAD engage in more frequent mutual chromatin contacts than do loci from different TADs. Consequently, TADs have been proposed to play a role in modulating transcription by promoting intra-TAD and preventing inter-TAD enhancer-promoter interactions, with potential implications in health and disease (*8–13*).

The functions of TADs and chromatin loops have also been investigated in the context of DNA replication (*14, 15*), V(D)J recombination (*16, 17*) and DNA repair, particularly, the repair of DNA double-strand breaks (DSB) (*18–21*). With respect to DSB repair, TAD boundaries appear to compartmentalize the chromatin response during the repair process, characterized by the mega-basepair scale formation of repair foci consisting of nucleosomes containing phosphorylated H2AX (γH2AX) and chromatin binding DSB repair proteins such as MDC1, 53BP1 and BRCA1 (*20, 21*). Specifically, ChIP-Seq measurements after a site-specific DSB in mammalian cells indicated that the extent of γH2AX, 53BP1 and MDC1 propagation correlates with the location of some TAD boundaries (*20, 21*). An explanation for the compartmentalization of the DSB repair chromatin response within individual TADs was suggested in (*21*). Based on the finding that DSBs induced by the AsiSI meganuclease can act as loops anchors, Arnould *et al* proposed that H2AX is phosphorylated by an ATM kinase located at the DSB, and that the propagation of the γH2AX domain is mediated by cohesin-dependent extrusion of a break-anchored loop (*21*). However, a recent work using live-cell imaging challenged this model by showing that cohesin subunits are recruited to DSBs significantly later than the recruitment of MDC1, the direct binding partner of γH2AX (*22*). Thus, formation of γH2AX-dependent DSB repair foci might occur independently of break-anchored loop extrusion. Interestingly, in yeast, site-specific DSBs have also been shown to accumulate cohesin (*23, 24*) and have been observed to act as loop anchors (*25*). However, in this case the loops were found to form during DSB repair by homologous recombination (HR) and in a DNA end-resection-dependent manner (*25*). Briefly, HR is a multistep pathway that entails: first, resection of each end of the DSB to generate long 3’ single-stranded (ss)DNA tails; second, loading of the RAD51 recombinase onto ssDNA to form the RAD51 nucleoprotein filament; third, the homology search—the process by which the RAD51 nucleoprotein filament searches for a homologous donor, culminating in RAD51-mediated strand exchange; fourth repair synthesis and termination of the HR reaction.

In summary, although several models have been proposed, the role of break-anchored chromatin loops in the processes of DSB repair remain poorly understood. Here, we investigated this question using a combination of time-course ChIP-Seq, time-course Hi-C and quantitative assays of DSB repair.

## RESULTS

### Cas9 breaks are anchors for cohesin loop extrusion

To study DSB-induced changes in the 3D genome, we induced DSBs using our recently developed multi-target CRISPR system, where a degenerate guide RNA (gRNA), or multi-target gRNA (mgRNA), directs a Cas9 nuclease to induce DSBs at >100 endogenous on-target sites (*26*). We nucleofected HEK293T cells with ribonucleoprotein complexes (RNPs) composed of purified Cas9 preassembled with the AluGG mgRNA, a gRNA we previously characterized (*26*), and performed *in situ* Hi-C (*2*) 3 h after delivery of the CRISPR/Cas9 ribonucleoprotein (RNP) as well as in control, untreated cells. For each condition, Hi-C was done in biological replicates and, after confirming a strong reproducibility (**Fig. S1**), replicates were merged totaling ~1B contacts per condition, which allowed us to build high-resolution Hi-C contact maps. Visual comparison of Hi-C maps around select Cas9 breaks revealed stripes that emanated from the predicted location of the Cas9 break (**Fig. 1a**). These stripes, which represent interactions between the break site and neighboring loci, were absent in the untreated control sample, suggesting that localized DSB-chromatin interactions arise *de novo* because of Cas9 activity.

**Figure 1.**
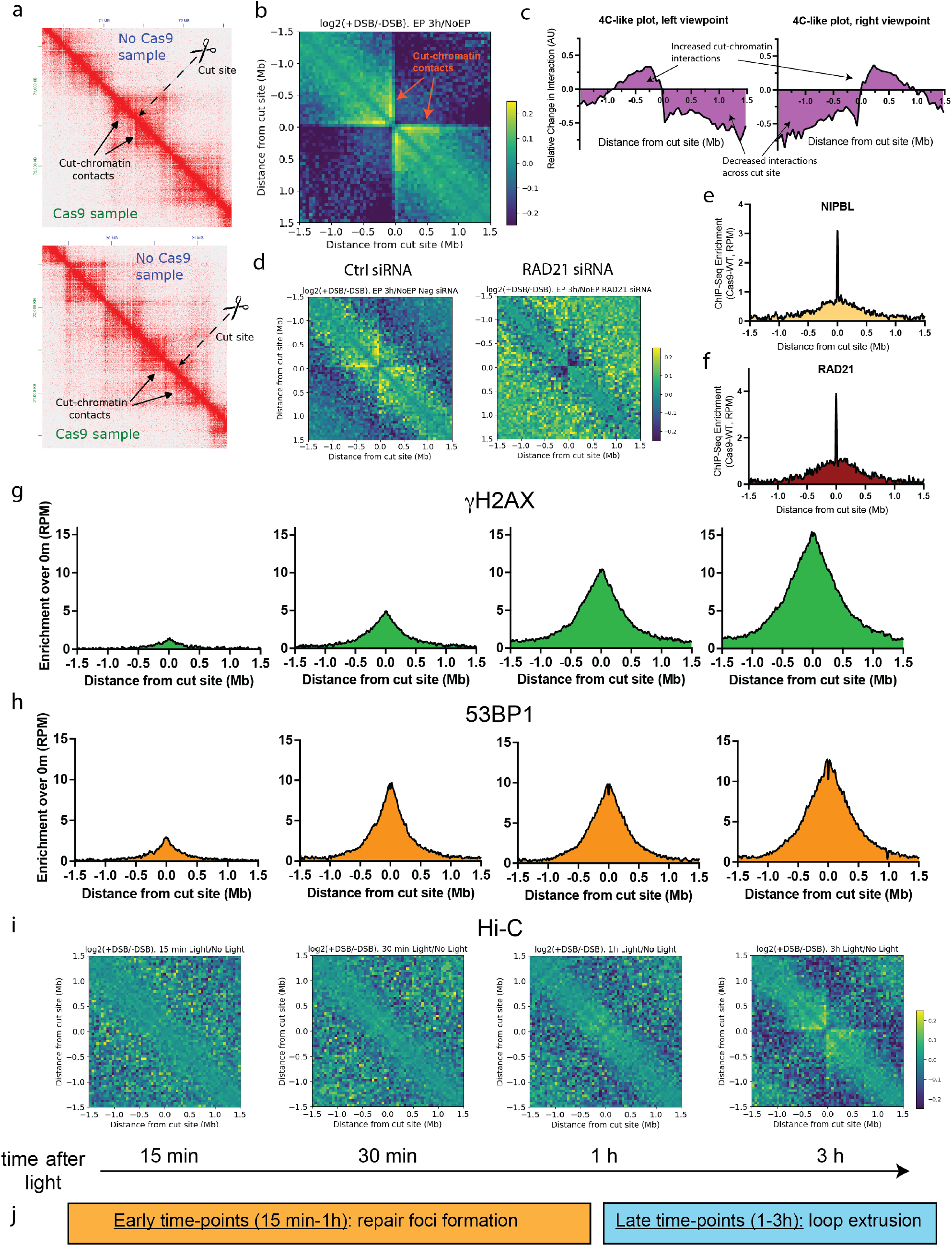
Break-anchored chromatin loops at Cas9 breaks. **a**, High-resolution (5 kb) Hi-C maps around two representative AluGG cut sites (in chr9, top; and chr14, bottom) in HEK293T. Untreated samples are shown in the top right triangle, Cas9-treated samples are shown in bottom left. **b**, Average log2 ratio of chromatin contacts around the best 100 AluGG cut sites with the strongest MRE11 enrichment. Hi-C matrices were retrieved at 50 kb resolution in a window of 3 Mb, centered at each MRE11 peak. **c**, 4C-like plots were computed from high-resolution Hi-C matrices around the top 100 MRE11 peaks using a left and right viewpoint with respect to the cut site for the Cas9-treated and undamaged sample. **d**, Cas9-induced chromatin contacts in control siRNA and RAD21 siRNA treated HEK293T cells. Differential Hi-C contacts were computed and plotted as in panel b. **e, f**, Cas9-induced NIPBL and RAD21 ChIP-Seq enrichment. ChIP-Seq profiles for NIPBL and RAD21 were averaged around the 126 AluGG on-target sites in Cas9-treated and untreated cells. The resulting untreated profile was subtracted from the Cas9-treated profile. **g-i**, Time-course γH2AX (g), 53BP1 (h) ChIP-Seq and Hi-C (i). HEK293T cells were treated with Cas9 RNP with caged AluGG gRNA for 12h, followed by light-induced Cas9 activation and harvest at 0 min, 15 min, 30 min, 1 h and 3 h. ChIP-Seq profiles were computed as described in panel **e, f**, and average Hi-C profile was obtained as in panel b, using the 0 min (no-light) control sample as reference. **j**, A proposed timeline.

To better visualize Cas9-induced changes in chromatin contacts, we computed the log2 of the ratio between the contacts in the Cas9 treated sample and the untreated sample averaged among the 100 best cleaved sites, as determined from previous MRE11 ChIP-Seq (*26*). The resulting differential Hi-C map revealed increased interactions between DSB sites and their neighboring chromatin, represented as cross-shaped yellow stripes (**Fig. 1b**), consistent with previous studies (*21, 27*) and with our inspection of individual cut sites (**Fig. 1a**). We further characterized these DSB-chromatin interactions by computing differential 4C-like plots using the regions adjacent to the breaks as viewpoints (**Fig. 1c**). DSB-chromatin interactions spanned a distance of ~1 Mb to the left and to the right of the cut-site but did not span the DSB locus itself. Rather, interactions between loci spanning the DSB site were diminished, suggesting that Cas9-induced DSBs acquire chromatin insulation properties, reminiscent of TAD boundaries. Thus, Cas9 breaks interact with their neighboring chromatin in an asymmetric fashion, and these interactions extend approximately 1 Mb from the DSB site.

Cohesin has been implicated in the formation of chromatin loops during repair of DSBs induced by the AsiSI meganuclease (*21*). To test whether cohesin is involved in the formation of DSB-chromatin interactions at Cas9-induced breaks, we delivered siRNA against RAD21 or a control siRNA into HEK293 cells, nucleofected the multi-target CRISPR RNP and performed Hi-C. RAD21 depletion abolished the DSB-chromatin contacts demonstrating that cohesin is instrumental in the formation of these structures (**Fig. 1d** and **Fig. S2**). Moreover, the loop extruding cohesin subunit NIPBL was recruited to Cas9 breaks, as determined by ChIP-Seq, as were two additional cohesin subunits, RAD21 and the DNA-damage phosphorylated form of SMC1 (pSMC1) (**Fig. 1e, 1f** and **Fig. S3**), further supporting a direct involvement of cohesin during Cas9 break repair. We conclude that Cas9-induced DSBs interact with their neighboring loci by a cohesin-driven mechanism, possibly involving loop extrusion.

### Break-anchored loops form after repair foci

Previous work reported the existence of break-anchored chromatin loops at meganuclease-induced DSBs, and the authors proposed that a cohesin-based loop extrusion mechanism is required for formation of the ~1 Mb γH2AX domain that flanks the DSB site (*21*). To test this hypothesis, we performed a time-course experiment using a light-activated, very fast (vf) CRISPR version of our multi-target system (*26, 28*). We measured the dynamics of repair foci formation via time-resolved ChIP-Seq against γH2AX and 53BP1 – two major factors known to decorate chromatin within the TAD that contains the DSB. In parallel, we performed time-resolved Hi-C to measure the time scale of break-anchored chromatin loop formation. We captured these events at 15 min, 30 min, 1 h and 3 h after CRISPR activation (**Fig. 1g-i**). γH2AX and 53BP1 were detected as early as 15 min after DSB induction (**Fig. 1g, h** and **Fig. S4**). The γH2AX signal spread over the entire time-course studied, with the largest changes observed in the early (15 min to 1 h) time points, whereas 53BP1 roughly saturated at 30 min. In contrast, our Hi-C data showed no break-anchored chromatin loops at the time points earlier than 1 h, and these loops only showed a robust signal at 3 h after damage (**Fig. 1i** and **Fig. S4**). These findings are consistent with recent live-cell imaging measurements showing that cohesin recruitment to a DSB occurs after the formation of MDC1 foci (*22*).

In summary, γH2AX and 53BP1 form a robust chromatin response to the DSB in less than 1 h, *well before* the formation of break-anchored loops, which occur between 1 h and 3 h after DNA damage (**Fig. 1j**). Thus, break-anchored loop extrusion is not a major driver of formation of the γH2AX domain.

### Break-anchored loops form during DSB processing for homologous recombination

If not for repair foci formation, is there a role for break-anchored chromatin loops in DSB repair? To address this question, we further analyzed our Hi-C data on a site-by-site basis to determine whether Cas9 sites with stronger changes in chromatin contacts are prone to undergo a particular repair pathway. In interphase nuclei, chromatin loops compartmentalize the genome into TADs; the insulation score metric quantifies TAD boundaries, with boundary locations showing a drop in insulation (*1, 29*). Using our high-resolution Hi-C data, we computed average insulation score profiles around the 126 AluGG gRNA on-target sites and observed a drop in insulation score in the Cas9-treated sample, but not the untreated one, consistent with Cas9 breaks acting as *de novo* TAD-like boundaries (**Fig. 2a**). A site-by-site analysis revealed a decrease in insulation score in 117 out of 126 on-target sites upon DNA damage, indicating that the observed change in average insulation score profiles does not stem from a few breaks, but rather, is a general DNA repair phenomenon (**Fig. 2b**).

**Figure 2.**
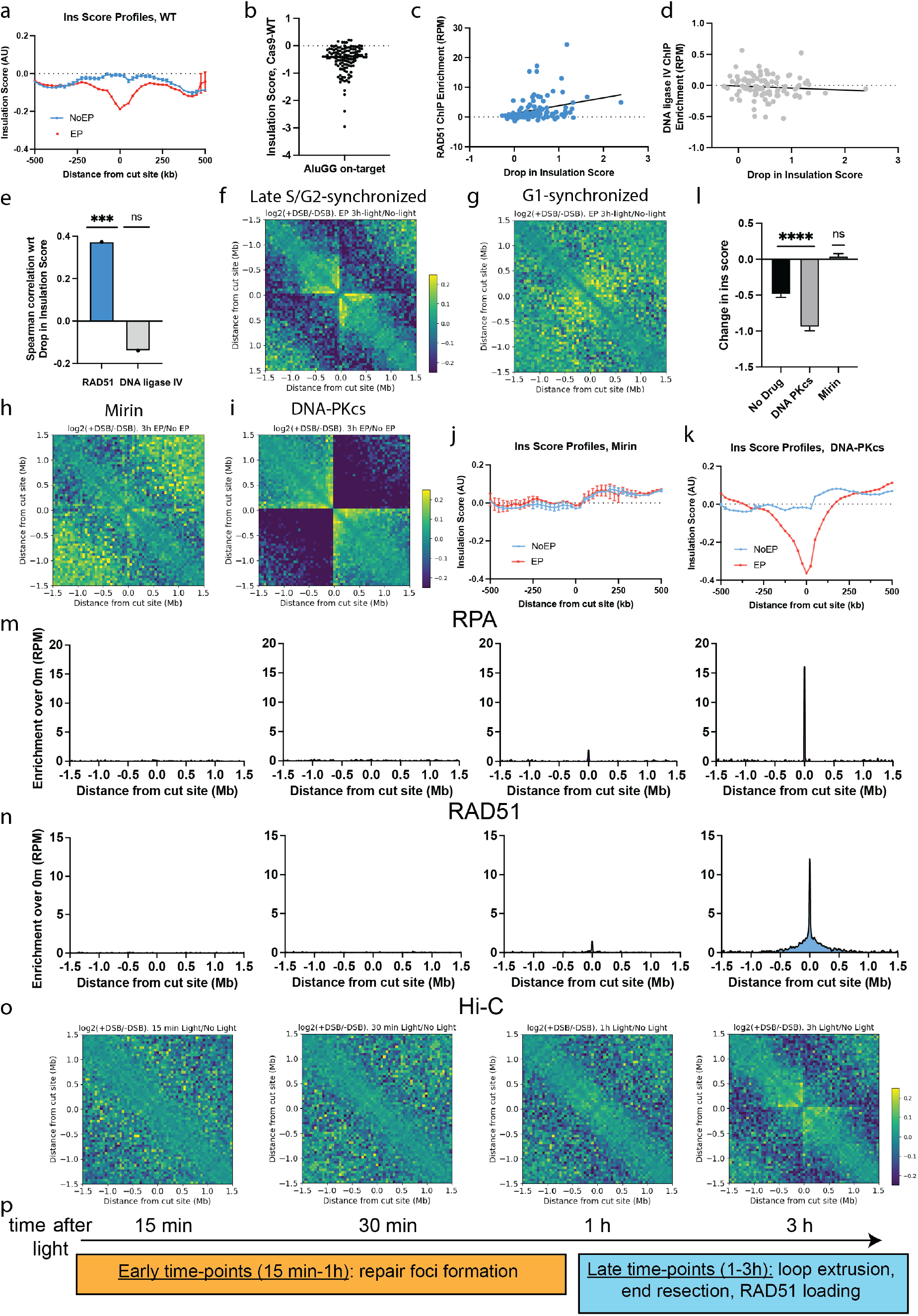
Break-anchored chromatin loops are formed during homologous recombination. **a**, Insulation score analysis around Cas9 breaks. Whole-chromosome Hi-C matrices were retrieved at 25 kb resolution and insulation score was then computed for each chromosome using the matrix2insulation script from the cworld package (*29*). Profiles were extracted in a window of 1 Mb around each of the 126 AluGG on-target sites and averaged. **b**, Change in insulation score upon Cas9 treatment was computed in a window of 50 kb centered around each AluGG on-target site. Note: reduced insulation score implies increased insulation. **c, d**, ChIP-Seq enrichment of RAD51 (c) and DNA ligase IV (d) vs drop in insulation score upon Cas9 treatment per cut site. Black lines are linear fits. **e**, Spearman correlation coefficient from (c) and (d). **f, g**, Averaged differential Hi-C contact maps in HEK293T cells synchronized in late S/G2 (f) or G1 (g). **h, i**, Averaged differential Hi-C contact maps treated with Mirin (h) or DNA-PKcs inhibitor (i). **j, k**, Insulation score profiles around on-target AluGG in cells treated with Mirin (j) or DNA-PKcs inhibitor (k). **l**, Cas9-induced change in insulation score around AluGG on-target sites in cells without drug, or treated with DNA-PKcs inhibitor or Mirin. Error bars are the standard error of the mean. Statistical significance with respect to null hypothesis was obtained using one sample t-test. **m-o**, Time-course RPA (m), RAD51 (n) ChIP-Seq and Hi-C (o). Hi-C plots are reproduced from **Fig. 1i** for each comparison. **p**, A proposed timeline.

To determine which repair pathway is dominant at each cut site, we performed ChIP-Seq for DNA ligase IV and RAD51, which have been used to classify DSBs as non-homologous end-joining (NHEJ) or homologous recombination (HR) prone, respectively (*30*). Consistent with both pathways being operative at Cas9 breaks, the average ChIP-Seq profiles revealed a damage-induced enrichment of both DNA ligase IV and RAD51 in the vicinity of Cas9 on-target sites (**Fig. S5**). We thus computed the Cas9-induced ChIP-Seq enrichment on a site-by-site basis and computed the Spearman correlation *r* between this ChIP-Seq enrichment and the drop in insulation score around the 100 best cleaved Cas9 DSB sites. RAD51 showed a significant positive correlation (*r*=0.37) with the damage-induced decrease in insulation score, whereas DNA ligase IV showed a weak, non-significant, negative correlation (*r*=-0.17) (**Fig. 2 c-e**). Thus, DNA breaks that are prone to undergo HR are associated with stronger insulation properties. Consistent with a role for cohesin in insulation properties at a DSB, all three cohesin subunits RAD21, pSMC1 and NIPBL, showed a significant positive correlation with the damage-induced drop in insulation (**Fig. S6**).

Based on these findings, we hypothesized that break-anchored loops form during processing of the DSB for repair by HR. To test this hypothesis, we implemented a protocol to induce Cas9 breaks at defined cell cycle stages, by exploiting our vfCRISPR system (*28*). In short, cells were synchronized using thymidine blocks and were then electroporated with a caged gRNA at the time of release from the block (**Fig. S7**). Cells were then exposed to 365 nm light to activate the caged vfCRISPR at defined time points following release from the block, corresponding to well-defined synchronized cell cycle stages (**Fig. S8, 9**). Using this approach, we activated vfCRISPR in cells synchronized in G1 and in late S/G2 phases – using, respectively, double and single thymidine blocks –, performed Hi-C 3h after DSB induction and compared the results with similarly synchronized, nucleofected cells that were not exposed to vfCRISPR-activating light. Cells synchronized in S/G2, which are able to perform HR, recapitulated the break-centered Hi-C stripes observed in unsynchronized cells, indicative of the formation of break-anchored loops (**Fig. 2f** and **Fig. S10**). Strikingly, G1-synchronized cells showed no appreciable change in chromosome contacts in response to Cas9-induced DSBs (**Fig. 2g** and **Fig. S10**), despite showing levels of DSB induction comparable to late S/G2 cells (**Fig. S11**). Given that HR is active in late S/G2 and inactive in G1, our results suggest that break-anchored loop formation occurs in relation to DSB processing for HR.

To further test the relationship between break-anchored loop formation and HR, we performed multi-target CRISPR delivery and Hi-C in the presence of a DNA-PKcs inhibitor, which enhances HR by inhibiting the catalytic activity of the non-homologous end joining kinase DNA PKcs (*31*), and mirin, a drug that abolishes HR by preventing MRE11 exonuclease-mediated DNA end resection (*32*). Notably, DNA-PKcs inhibition resulted in stronger break-anchored loops as apparent from brighter stripes, whereas treatment with mirin abolished the stripes (**Fig. 2h, i** and **Fig. S12**). As an additional quantification of the effect of these drugs in Cas9-induced chromosome conformation changes, we performed deeper sequencing of the Hi-C samples and conducted insulation score analysis (**Fig. 2j-l**). DNA-PKcs inhibition further accentuated the Cas9-induced drop in insulation score, suggestive of stronger loops, whereas mirin treatment eliminated insulation score changes upon Cas9 breaks. Therefore, we conclude that break-anchored cohesin loops are formed during HR in a DNA end resection-dependent manner.

### Time-course measurements support a role for break-anchored loop extrusion in HR

We investigated the temporal relationships between DNA end resection, RAD51 loading and break-anchored loop extrusion, using our light-activated multi-target vfCRISPR. To this end, we performed a second time-course experiment, comparing the accumulation of signals for RPA and RAD51 (by use of ChIP-seq) at vfCRISPR-induced DSBs with the formation of break-anchored loops (by use of Hi-C) at the same sites (the latter from **Fig. 1i**).

RPA and RAD51 were undetectable within the first 30 min after DSB induction, revealing a strong, Cas9-induced enrichment only at 3 h (**Fig. 2m, n** and **Fig. S4**). Thus, in our multi-target vfCRISPR system, resection and RAD51 loading occur at relatively late time points (> 1 h). Strikingly, this delay in RPA and RAD51 recruitment matched closely the dynamics of loop formation captured by time-resolved Hi-C (**Fig. 2o** and **Fig. S4**), further supporting a causal link between break-anchored loops and HR.

In conclusion, our time-resolved data revealed two different time-scales during Cas9-induced DSB repair (**Fig. 2p**): first, γH2AX and 53BP1 chromatin responses are detectable at early time points (15 min – 1 h), whereas DNA end resection, RAD51 loading, and break-anchored loop formation occur at late time points (1-3 h).

### A broad, RAD51 chromatin domain reflects the homology search

Our time-course experiment yielded another unexpected finding. Whereas the RPA ChIP-seq signal at 3 h was constrained to a narrow window (~5 kb) centered at the cut site, the RAD51 profile propagated to long distances of several hundreds of kilobases away from the break. To better quantify this phenomenon, we performed ChIP-seq for RPA and RAD51 3 h after delivery of regular, non-caged AluGG gRNA. The resulting RPA ChIP-Seq profile showed a peak spanning a width of 4.8 ± 0.7 kb, centered at the cut site (**Fig. 3a, b**), a finding that is consistent with the time-course data and with previous reports showing that DNA end resection extends for a few kilobases from the DSB (*33*). The RAD51 profile showed a peak of similar width of 6.1 ± 0.7 kb, which we attribute to ssDNA that is coated by the recombinase (**Fig. 3c, d**). Yet, in addition to this peak, the average RAD51 ChIP-Seq signal spread to a broad domain of 600 ± 30 kb in size (**Fig. 3d, e**), consistent with our time-course measurements (**Fig. 2n**). Visual inspection of RPA and RAD51 ChIP-Seq signals at individual AluGG cut sites corroborated the presence of these broad RAD51 domains that extend beyond the main, narrow peak (**Fig. S13**). An alternative RAD51 antibody that binds a different epitope gave a similar average ChIP-Seq profile (**Fig. S14**) with indistinguishable widths for both the narrow peak (P=0.85) and the broad RAD51 domain (P=0.54) (**Fig. 3e**).

**Figure 3.**
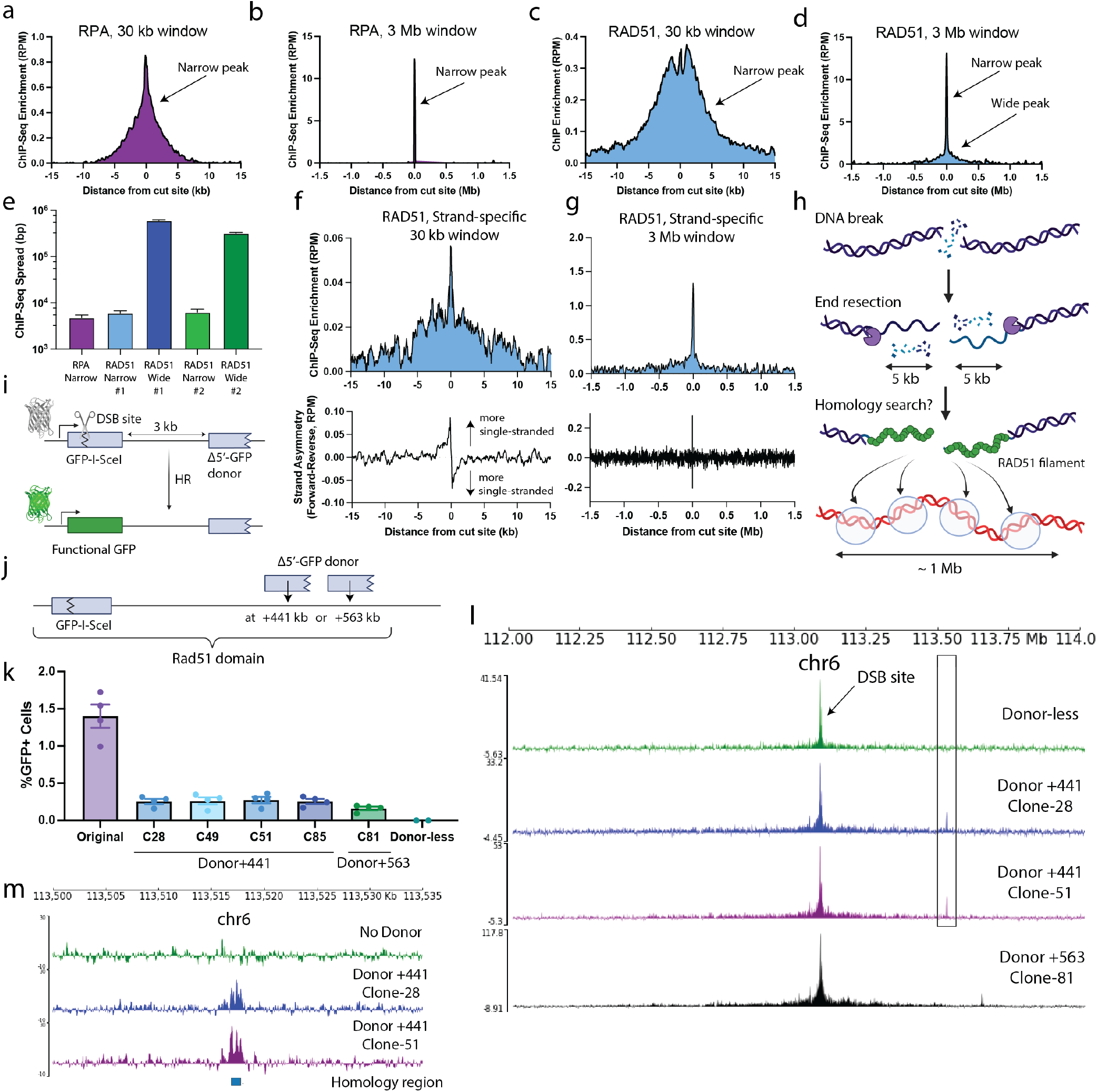
Broad RAD51 ChIP-Seq profiles inform on homology search. **a-d**, Average RPA (a, b) and RAD51 (c, d) ChIP-Seq profiles around AluGG on-target sites in a window of 30 kb (a, c) and 3 Mb (b, d). Arrows mark narrow (RPA, RAD51) and broad (RAD51) peaks. **c**, Average width of RPA and RAD51 peaks. Error = standard error of the mean. **f, g**, Strand specific ChIP-Seq of RAD51. Top panels are regular ChIP profiles (30 kb and 1 Mb windows). Bottom panels show the strand asymmetry, computed as the difference between the forward and the reverse reads. **h**, Cartoon depicting the interpretation of the broad RAD51 profile as a measure of homology search. **i**, Schematic of the HR-GFP reporter system to measure DSB-induced HR. *GFP-I-SceI heteroallele* harbors a site for DSB induction. HR repair via use of a 5’-Truncated GFP donor (~3 kb away from the DSB) generates WT GFP that can be measured in flow cytometry. **j**, Schematic of the reporter system designed to study HR using distant donors within the TAD. A monoclonal mES cell clone was generated with a single *GFP-I-SceI* copy at Rosa26 on Chr6. Then, derivative monoclonal lines were made with the *Δ5’-GFP* HR donor at +441 kb or +563 kb from the *GFP-I-SceI* copy. **k**, HR readout for the original cell line (with the donor at ~3 kb from the break), four Donor+441 clones, a Donor+563 clone, and the donor-less clone. **l**, RAD51 ChIP-Seq profile in donor-less cells (top), in two of the Donor+441 clones and the Donor+563 clone. **m**, Zoom-in of (l) showing the RAD51 signal over *Δ5’-GFP* in the donor-less clone vs. two Donor+441 clones. The donor-less clone ChIP-Seq data in panels m, l was aligned to a genome containing a +441kb donor.

RAD51 binds both ssDNA and dsDNA (*34*). We thus hypothesized that the broad RAD51 domain, which lacks RPA binding, represents RAD51-dsDNA interactions. To test this, we induced DNA breaks using AluGG gRNA and measured RAD51-chromatin binding at 3 h using strand-specific ChIP-seq, a method that reveals the degree of strand asymmetry in the ChIP sample, with higher strand asymmetry corresponding to a higher proportion of ssDNA molecules of a specific polarity (*35*). The RAD51 strand-specific ChIP-Seq profile showed similar qualitative features to the regular ChIP-Seq profile, including a peak at the cut site and a broad chromatin domain (**Fig. 3f, g** and **Fig. S15**). Analysis of strand asymmetry at each chromosomal location revealed that the extent of RAD51-ssDNA binding is constrained to a region of ~2 kb centered at the break, with the expected switch in polarity on either side of the DSB, reflecting RAD51 loading onto 3’ ssDNA DNA ends (*35*). In contrast, at distances >2.5 kb from the DSB, strand asymmetry was lost, implying a RAD51-dsDNA binding mode.

Studies *in vitro* suggest that, during the homology search, the Rad51-ssDNA nucleoprotein filament interacts with a dsDNA template and scans it locally through one-dimensional random walk, as part of the homology search (*36*). If the dsDNA is homologous to the RAD51 nucleoprotein filament, HR proceeds with formation of a D-loop. In contrast, the RAD51 nucleoprotein filament will dissociate from a non-homologous dsDNA (*36*). In yeast, Rad51 ChIP-Seq after a site-specific DSB was suggested to reflect the extent of the homology search, with the Rad51 ChIP-Seq profile spanning a broad chromatin domain that represents Rad51-dsDNA interactions during chromatin scanning (*37*).

We asked whether – similar to the yeast case – the broad RAD51 domains that we observed in mammalian cells have features characteristic of homology search (**Fig. 3h**), in particular, whether it shows local accumulation at an appropriate HR donor (*36, 37*). We adapted a previously described HR reporter, which contains a *GFP* heteroallele (*GFP-I-SceI*) that entails a full length *GFP* gene that is interrupted by a target site for the endonuclease I-SceI, and a 5’ truncated (*Δ5’-*) *GFP* recombination donor situated ~3 kb from the *I-SceI* site (**Fig. 3i**) (*38*). DSB-induced HR is thus quantified by the production of GFP^+^ cells. Using a monoclonal mouse embryonic stem (mES) cell line that contains a single-copy HR reporter at the *Rosa26* locus of Chr6, we first removed the *Δ5’-GFP* donor *via* a CRISPR/Cas9-induced deletion. We nucleofected a Cas9-gRNA RNP targeted to the *I-SceI* site to induce a DSB within *GFP-I-SceI* in these “donor-less” cells, and measured the RAD51 domain flanking this DSB using Rad51 ChIP-seq. The broad RAD51 domain covered chromosomal regions extending approximately −800 kb to +700 kb around the break site (**Fig. S16**). To determine whether the Rad51 signal is enriched over the HR donor during a successful homology search, we generated a new *Δ5’-GFP* donor containing select silent point mutations (conserving the amino acid sequence of GFP) sufficient to distinguish it from *GFP-I-SceI* during ChIP-seq sequence alignment. We targeted the new *Δ5’-GFP* donor to positions +441 kb (“Donor+441”) or +563 kb (“Donor+563”) relative to the *Rosa26*-located *GFP-I-SceI* (**Fig. 3j**), generating four independent Donor+441 clones and one Donor+563 clone in which a single, intact *Δ5’-GFP* donor was targeted to the same copy of Chr6 as the *Rosa26*-located *GFP-I-SceI* heteroallele (see Methods). *I-SceI*-induced HR in Donor+441 clones and the Donor +563 clone was readily detected but was less efficient than in the original HR reporter cell line (**Fig. 3k**; see Methods). Consistently, the donor-less clone showed no detectable HR levels (**Fig. 3k**). We then used Cas9-gRNA RNP nucleofection to induce a DSB at *GFP-I-SceI* in two Donor+441 clones and the Donor+563 clone and performed RAD51 ChIP-Seq 3 h after Cas9 delivery. Of note, in addition to the above-noted distributions of RAD51 ChIP-seq signal, the broad RAD51 peak contained an additional, narrow and more intense peak at the location of the HR donor, at either +441 kb or +563 kb (**Fig. 3l** and **Fig. S17**). This enhanced RAD51 signal over the *Δ5’-GFP* HR donor likely represents the result of a successful homology search by the RAD51 nucleoprotein filament(s) derived from the DSB at *GFP-I-SceI*. A zoomed-in view revealed that this RAD51 peak falls at the exact location of the *Δ5’-GFP* donor and shows three subpeaks, one on one end and the other two on the other end of this region (**Fig. 3m**). To exclude the possibility that the RAD51 ChIP-Seq peak at the *Δ5’-GFP* donor might be a read alignment artifact due to the high sequence similarity between the *Δ5’-GFP* donor and *GFP-I-SceI*, we aligned the RAD51 ChIP-Seq data derived from the “donor-less” HR reporter clone to a reference genome that includes the *Δ5’-GFP* donor (at positions +441 or +563) and observed no specific Rad51 enrichment over the *Δ5’-GFP* donor (**Fig. 3l, m** and **Fig. S17**). Thus, RAD51 ChIP-seq reads that faithfully map to the *GFP-I-SceI* heteroallele at *Rosa26* are not prone to artifactual alignment with the *Δ5’-GFP* heteroallele. In conclusion, the broad RAD51 domain shows local enrichment over the *Δ5’-GFP* HR donor, likely as a result of a successful homology search.

### Loop extruding cohesin promotes RAD51-chromatin interactions within the damaged TAD

Our time-course data from **Fig. 2** indicated that the formation of the broad RAD51 domain occurs contemporaneously with the break-anchored chromatin loops, suggesting that the break-anchored loops might participate in the homology search. We thus evaluated the effect of TAD boundaries on the spread of the broad RAD51 ChIP-Seq signal obtained in HEK293T subjected to AluGG DSBs (data from **Fig. 3c**). We computed the averaged, RAD51 ChIP-Seq enrichment around TAD boundaries that lie between 200 kb and 700 kb away from the 100 most efficiently cleaved AluGG cut sites (according to MRE11 ChIP-Seq) and noted a sharp decrease in RAD51 signal at the exact location of the TAD boundary (**Fig. 4a, b**). As control, we performed the same analysis around random, non-TAD boundary sites and observed no significant reduction in RAD51 signal (**Fig. 4b**). Therefore, at least some TAD boundaries correlate with the extent of the broad RAD51 peak, possibly restricting the homology search to the damaged TAD. To explore this idea at the level of a single locus, we treated HEK293T cells with a Cas9 complexed with an ACTB-targeting gRNA, previously shown to trigger Cas9 cleavage with high efficiency (*28*), and performed RAD51 ChIP-Seq. The resulting profile showed a drop at the nearest TAD boundaries (boundaries −1 and +1) and reached background levels upon crossing the next-to-nearest boundaries (−2 and +2 from the cut) (**Fig. 4c**). Similar findings regarding the drop in RAD51 signal at TAD boundaries were obtained using an additional gRNA targeting the MYC gene (**Fig. S18**).

**Figure 4.**
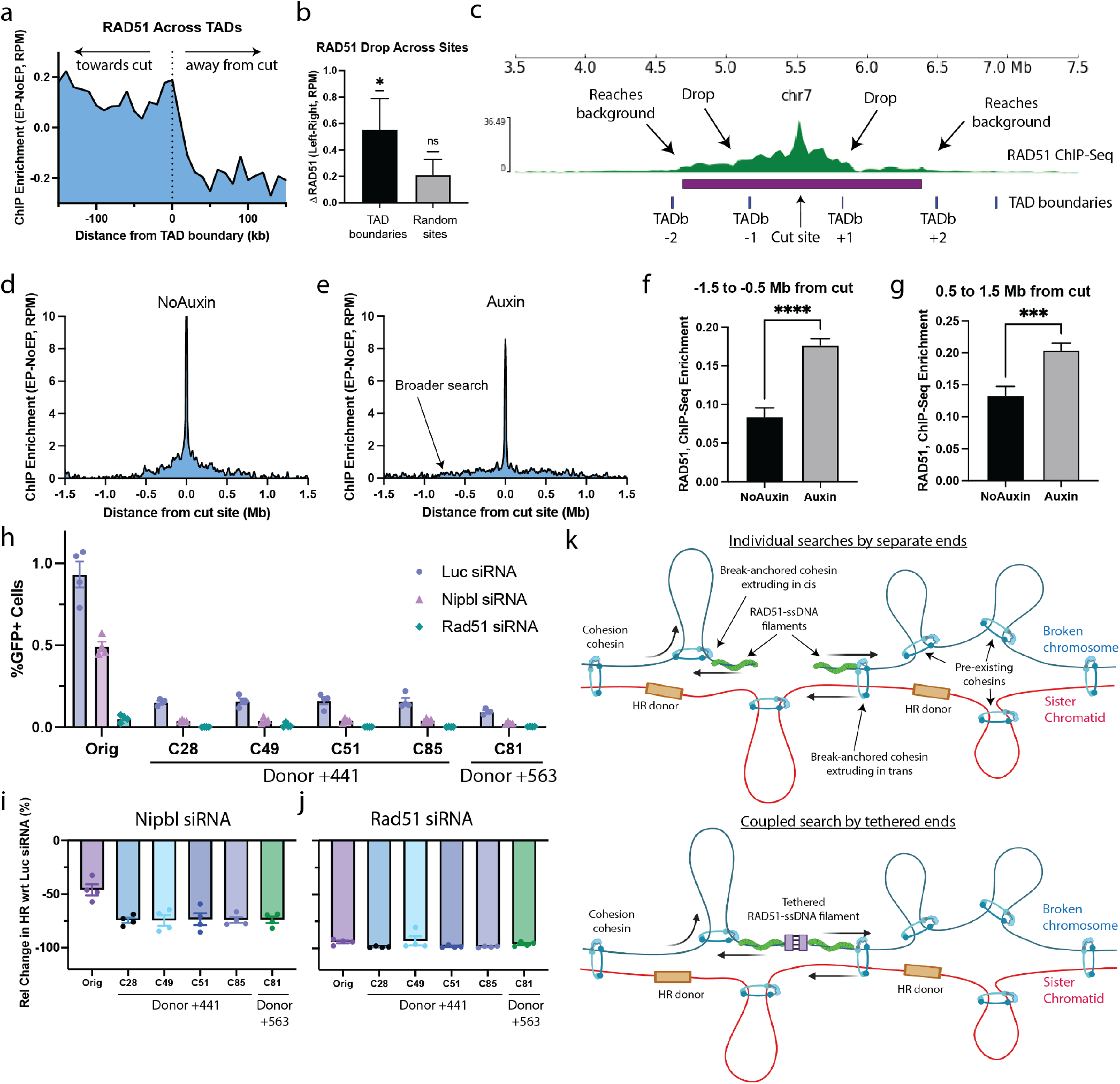
Loop-extruding cohesin regulates homology search and overall HR efficiency. **a**, RAD51 ChIP-Seq enrichment averaged around all TAD boundaries that lie 200 kb to 700 kb away from the 100 best Cas9 cut sites. **b**, RAD51 drop at TAD boundaries and random sites (picked at distances between 200 kb and 700 kb away from cut sites). Error = standard error of the mean. One-sample t-test was done to quantify significance from null hypothesis. **c**, RAD51 ChIP-Seq profile at ACTB gene in HEK293T. **d, e**, Average RAD51 ChIP-Seq profiles after Cas9-AluGG-gRNA-induced DSBs in HCT-WAPL-AID2 cells without (d) or with (e) auxin. **f, g**, Average change in RAD51 ChIP-Seq enrichment was obtained at regions −1.5 to 0.5 Mb (f) or 0.5 to 1.5 Mb (g) away from the cut. Error = the standard error of the mean. Unpaired t-test was done to quantify significance between the two conditions. **h**, HR-GFP assays were performed on the monoclonal cell lines from Fig. 3k after treatment with siRNA against luciferase (Luc), Nipbl or Rad51. **i, j**, Quantification of h, showing changes in HR following depletion of Nipbl-(i) or Rad51-induced (j), normalized to Luciferase siRNA HR levels. **k**, Cartoon depicting different mechanisms for cohesin-driven chromatin scanning during homology search.

Next, we tested the role of chromatin loops on the extent of the broad RAD51 domain. WAPL unloads cohesin from chromatin and acute WAPL degradation results in elongated loops, likely by extending cohesin’s residence time on chromatin (*39*). To manipulate WAPL levels, we employed a previously characterized HCT116 cell line, hereafter HCT116-WAPL-AID2, where WAPL is endogenously tagged with a degron system that permits fast, acute degradation upon the addition of auxin with minimal leakage (*15, 40*). We delivered multi-target CRISPR into the HCT116-WAPL-AID2 and compared the extent of spread of the broad RAD51 peak in the presence and absence of auxin. Consistent with a role of loop extrusion in the homology search, we found that WAPL loss resulted in a broader RAD51 ChIP-Seq domain (**Fig. 4d, e** and **Fig. S19**). Specifically, subtraction of the RAD51 profile in the absence (Auxin) and presence (No-Auxin) of WAPL revealed that regions that are further than 0.5 Mb from the DSB site have significantly higher levels of RAD51 after WAPL degradation (**Fig. 4f, g** and **Fig. S19**). These findings suggest that elongated chromatin loops result in a broader homology search.

Our observations so far suggest that loop-extruding cohesin could play a role in HR by promoting the homology search within the TAD that contains the DSB. To further explore the potential role of loop-extruding cohesin in HR, we studied the impact of NIPBL depletion on DSB-induced HR. NIPBL is a cohesin subunit that is required for loop extrusion but not for sister chromatid cohesion; consequently, NIPBL knock-down perturbs the function of cohesin loops without disrupting sister chromatid cohesion (*41*). We co-transfected mES cells containing the original HR-GFP reporter with an *I-SceI* expression vector together with control, *Nipbl* or *Rad51* siRNA and determined HR efficiency 72 h later (see Methods). We repeated the same procedure in four independent Donor+441 HR clones, the independent Donor+563 HR clone and the “donor-less” clone described above (**Fig. 4h**). As expected, no HR products were detected in the “donor-less” clone (**Fig. S20**). Depletion of NIPBL in cells containing the original HR reporter reduced HR efficiency by ~46% in comparison to control siRNA (**Fig. 4i**). This finding suggests that cohesin-mediated loop extrusion contributes to DSB-induced HR, even when the donor is located only ~3 kb from the DSB. Of note, the impact of NIPBL depletion was more pronounced in the Donor+441 and Donor+563 cell lines, where the HR donor is located far from the DSB site. NIPBL depletion resulted in mean HR reductions of 74% in Donor+441 clones, and 74% in the Donor+563 clone (**Fig. 4i**). Thus, at large DSB-donor distances, the dependence of HR on NIPBL becomes larger. As opposed to NIPBL, RAD51 depletion had impacts on HR that were independent of the DSB-donor distance, with reductions of 94% (original HR reporter), 97% (average over Donor+441 clones) and 96% (the Donor+563 clone; **Fig. 4j**). Taken together, our data suggests that loop extruding cohesin promotes HR by facilitating the homology search within the local TAD. Of note, the dependence of HR on loop extrusion is more pronounced when the homologous donor is positioned at a large distance from the DSB.

### A role for cohesin loop extrusion in the homology search

Based on our data, we propose that loop-extruding cohesin facilitates the homology search via directed 1D scanning of the local TAD. In post-replicative cells, DNA breaks can be repaired by HR through the assembly of a RAD51 filament at each resected DNA end (*42*). This filament can then perform a local search on the sister chromatid, which is discontinuously tethered to the broken chromosome by pre-existing deposits of cohesive cohesin (*41*). If the HR donor is not within reach of this local search, possibly because sister chromatids might not be perfectly aligned, or if the sister chromatid were also broken so that an alternative donor needs to be used for HR, this initial local search would be unproductive. Cohesin recruitment to the break site could then facilitate extrusion of a chromatin loop anchored at the break, allowing the RAD51 filament to travel along the chromosome until a new location is reached (see **Fig. 4k**). This filament would then perform a local search at this new location, increasing the chances of finding an appropriate HR donor.

Our findings invite investigation about additional aspects of homology search. For example, is cohesin extruding a loop *in cis* on the broken chromosome or is it extruding the sister chromatid by the broken end *in trans*, directionally scanning one DNA molecule with respect to the other (**see Fig. 4k**)? Another open question is whether the two ends of the DSB remain tethered to one another during the homology search, or whether they each perform a separate individual search (compare top and bottom panel in **Fig. 4k**). In yeast, the Rad51 nucleoprotein filaments on both sides of the break were proposed to be held together during the homology search, suggesting a coupled scanning mechanism (*25*), however this model remains to be tested in mammalian cells. How dynamic is the search and the formation of break-anchored chromatin loops? It has been shown that chromatin loops at the mouse Fbn2 TAD are transient and rare (*43*). It is likely that the break-anchored loops formed during HR are also transient, effectively increasing the 1D mobility of the break, rather than stably positioning it at a defined genomic location. In addition, the role in homology search of factors other than cohesin remains to be investigated. Of particular relevance is the HR factor Rad54, a key player in the late steps of HR, which has been shown to drive 1D scanning of the Rad51 filament along a stretch of DNA in vitro (*36, 44*).

The homology search problem posits the question: how does a broken DNA find a correct HR template in the mammalian genome? A 3D exploration mechanism, e.g. via 3D diffusion, has been proposed to drive homology searches during alternative lengthening of telomeres repair of telomeric DSBs (*45*). However, in non-repetitive regions, a 3D search mechanism is likely to be slow and inefficient (*46*). Dimensionality reduction via 1D local scanning can significantly accelerate this process, for example through sliding of the recombinase-ssDNA filament along dsDNA (*47*). However, because the two sister chromatids are only loosely connected, such scanning would require a highly processive and fast motor. Our data suggests that cohesin can facilitate this long-range scanning via loop extrusion. In the bacterium *Caulobacter crescentus*, live-cell imaging measurements revealed a fast, directional movement of bacterial recombinase (RecA) filaments over the entire bacterial chromosome on the order of minutes (*48*). This movement required another SMC protein, RecN, and the authors suggested that RecN could actively drive the search via 1D scanning. In yeast, chromatin loops are formed during HR, promoting break-chromatin interactions and facilitating contact with an HR donor *in cis* (*25*). Our data suggests that cohesin also plays a role in the homology search in mammalian cells, likely by promoting repeated long-range scanning of RAD51-ssDNA filaments against the sister chromatid. Taken together, these findings point to a universal role for SMC proteins in driving a directed 1D scanning during the homology search, likely conserved across different domains of life.

## Acknowledgments

This article is subject to HHMI’s Open Access to Publications policy. HHMI lab heads have previously granted a nonexclusive CC BY 4.0 license to the public and a sublicensable license to HHMI in their research articles. Pursuant to those licenses, the author-accepted manuscript of this article can be made freely available under a CC BY 4.0 license immediately upon publication. The authors would like to thank Claudia Carcamo, Sergei Rudnizky, and Reza Kalhor for stimulating discussions throughout the project. We are grateful to the GRCF and single-cell and transcriptomics cores for NextSeq and NovaSeq experiments and to Winston Timp and Loyal Goff at Johns Hopkins University for granting us access to their MiSeq machines. We are also grateful to Andrew Feinberg (Johns Hopkins) and the biopolymers core (HMS) for access to covaris sonicators.

## Author contributions

A.M.-G. and A.T.R. performed ChIP-Seq and Cas9 RNP nucleofections. A.M.-G performed Hi-C experiments. A.M.-G. analyzed ChIP-Seq, strand-specific ChIP-Seq and Hi-C data. N.M.N. generated and validated clonal mES cell lines with help from A.M.-G. and D.N. N.M.N. conducted HR-GFP assays. V.K.-N. and A.M.-G. validated Cas9 RNP nucleofections in mES cells. A.G.L. performed strand-specific ChIP-Seq. A.T.R., N.M.N. and D.N. performed Western Blots for siRNA validation. R.S.Z. assisted with data analysis. F.A.-V. assisted in initial conceptualization. M.T.K. provided WAPL-AID cell lines. A.M.-G. prepared figures and wrote the manuscript with input from T.H., R.S, A.T.R and N.M.N. T.H. and R.S. supervised the project.

## Data and materials availability

All data associated with this study are present in the main text or the supplementary materials. ChIP-seq and Hi-C data have been uploaded to the Sequence Read Archive under BioProject accession PRJNA1214218. Analysis code is available on GitHub (https://github.com/AlbertoMarinG/HomSearch/).

